## Supplementary figures and images for "Whole-genome analysis of Nigerian patients with breast cancer reveals ethnic-driven somatic evolution and distinct genomic subtypes"

### Supplementary Fig. S1

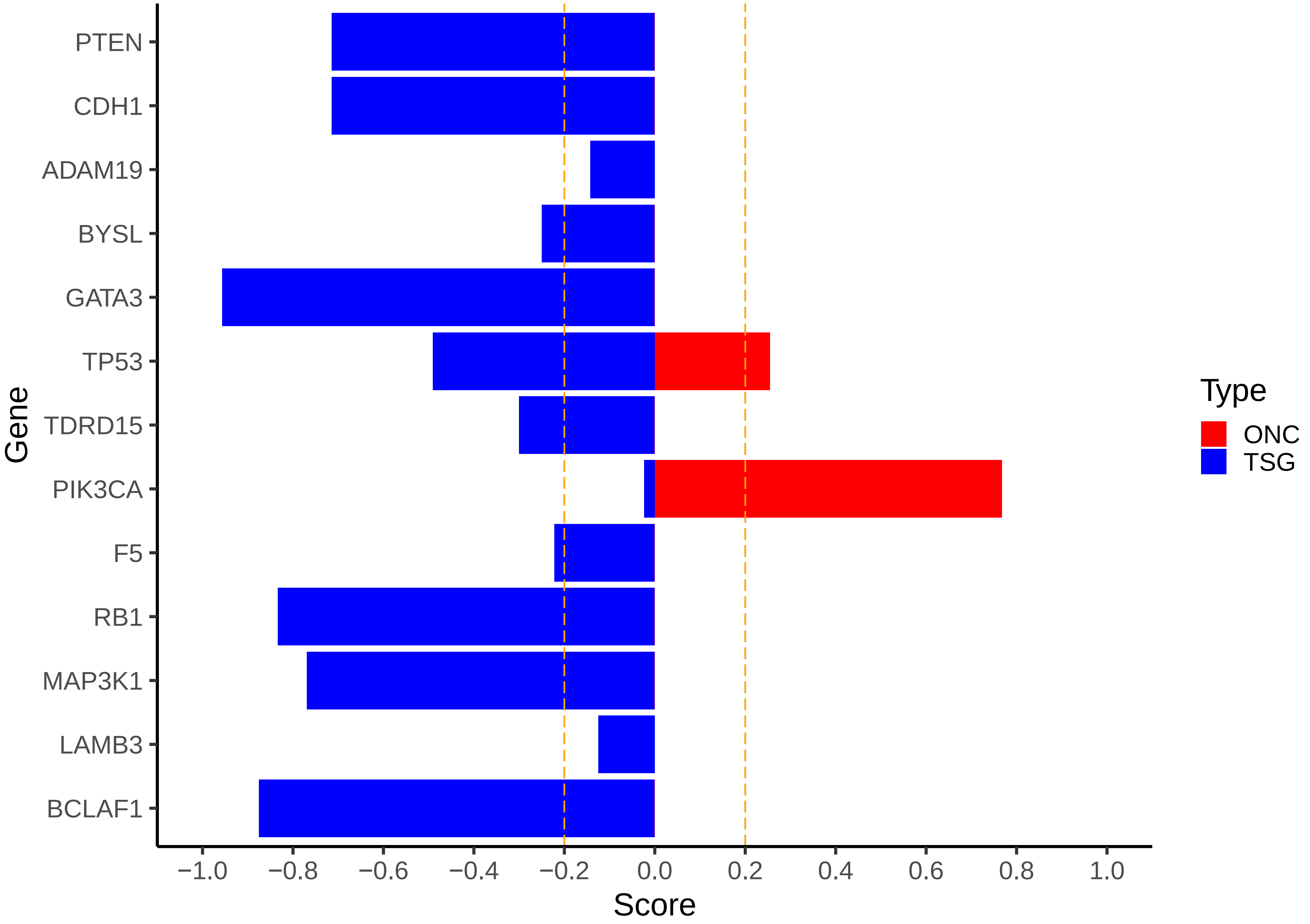

### Supplementary Fig. S2

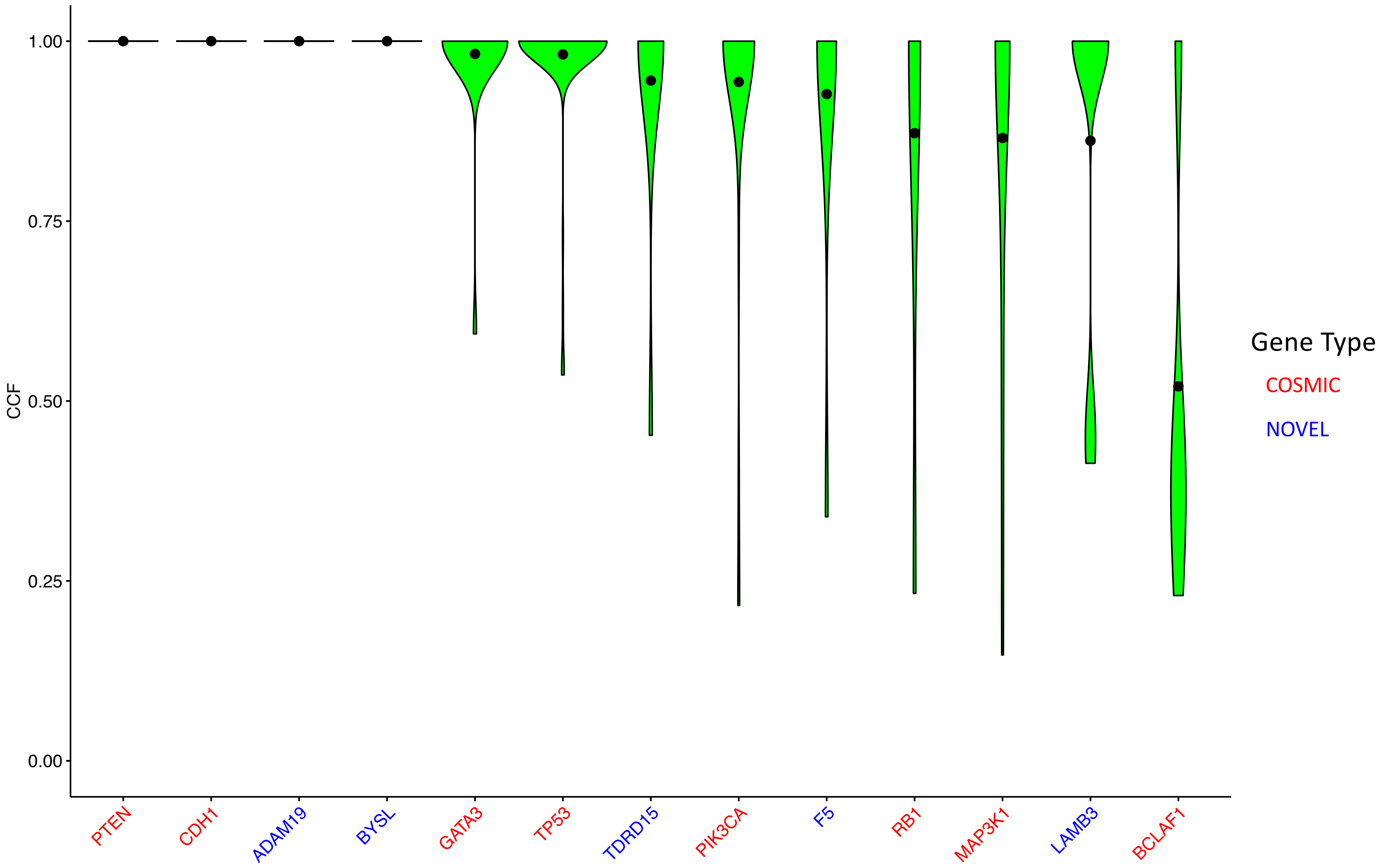

### Supplementary Fig. S3

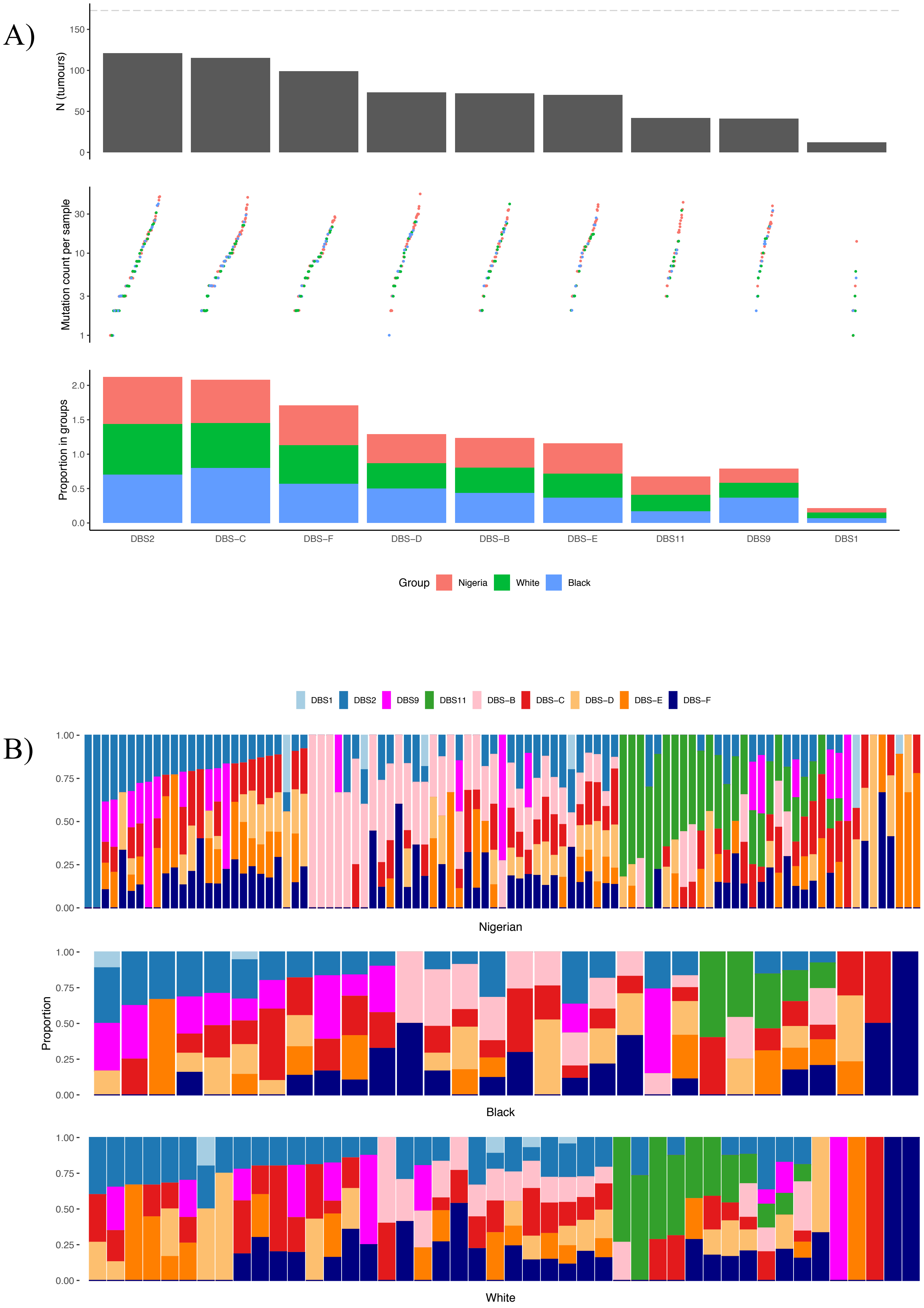

### Supplementary Fig. S4

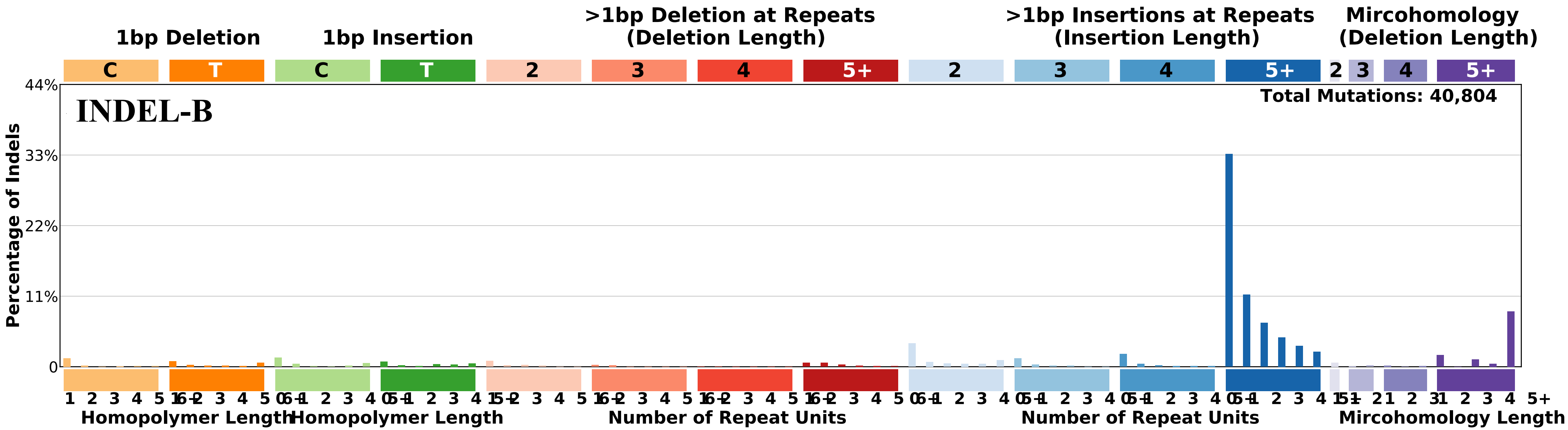

### Supplementary Fig. S5

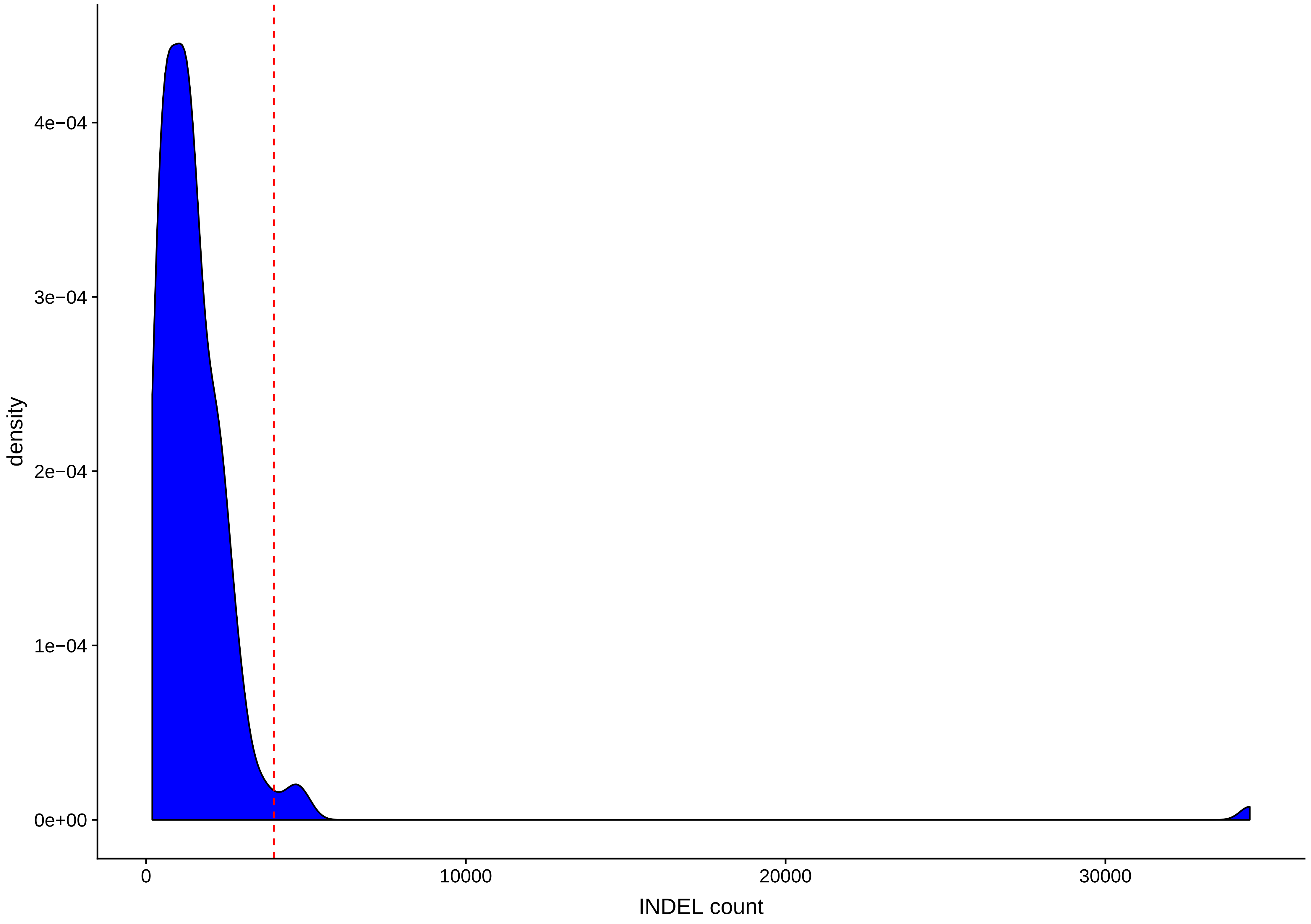

### Supplementary Fig. S6

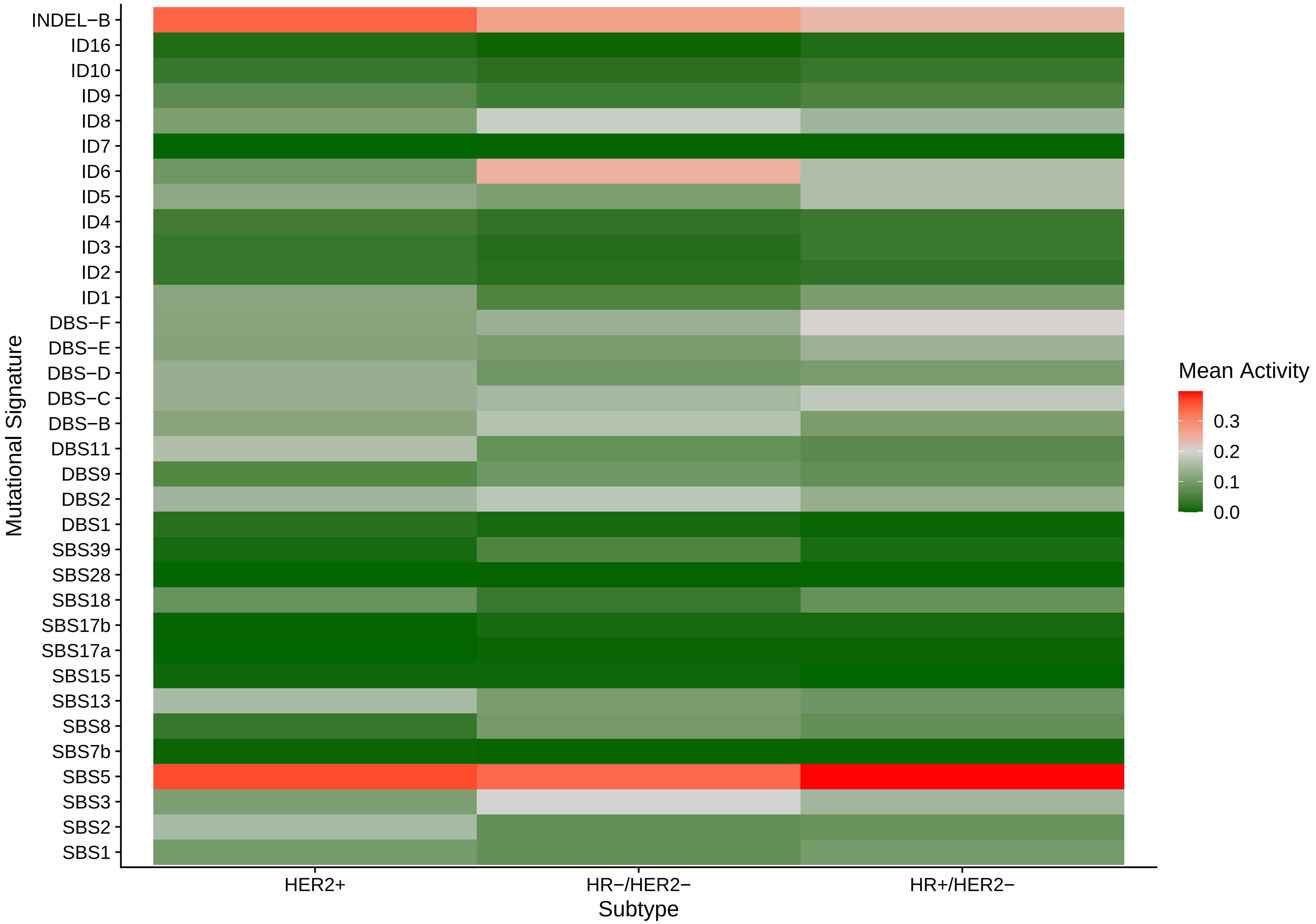

### Supplementary Fig. S7

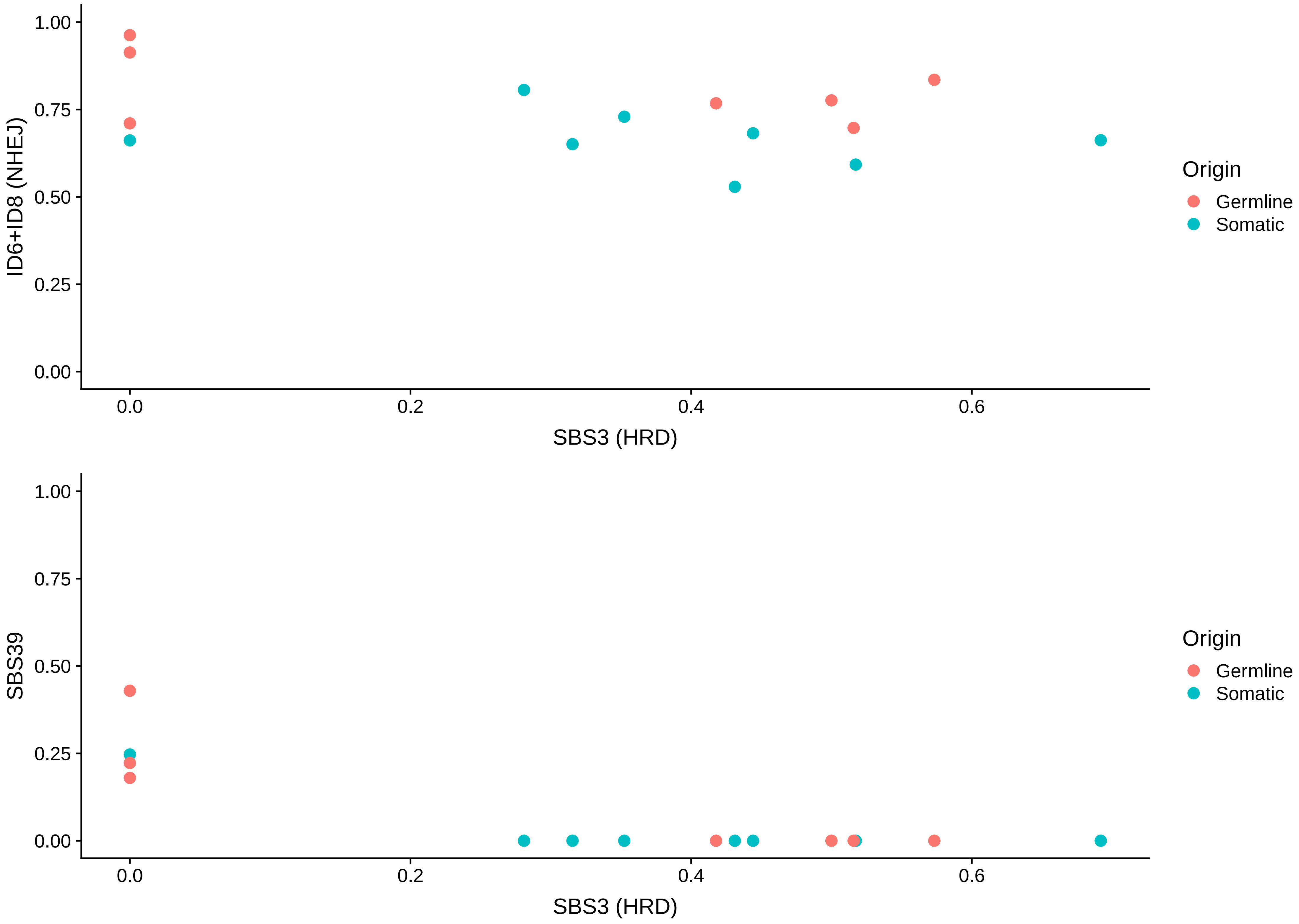

### Supplementary Fig. S8

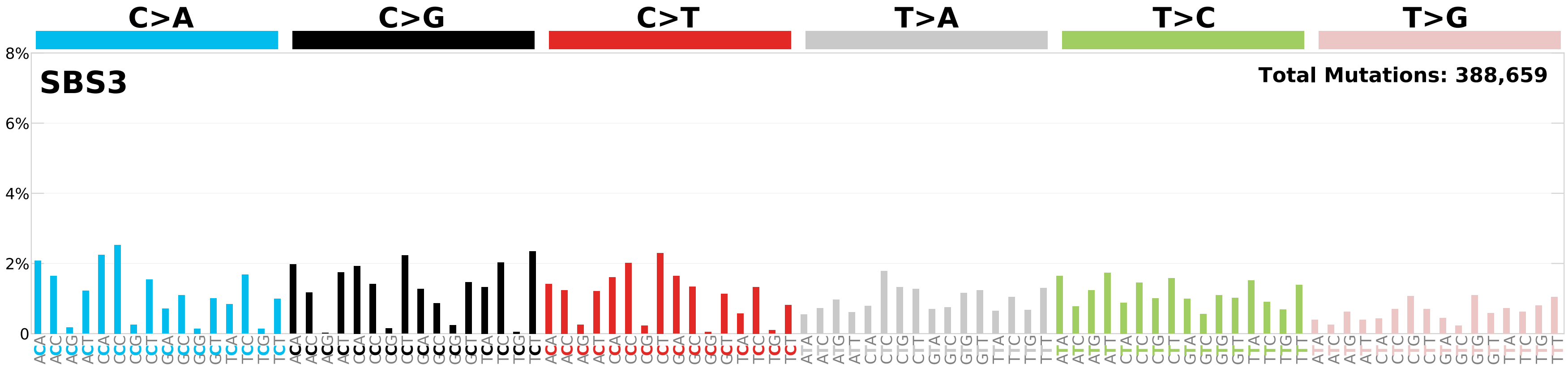

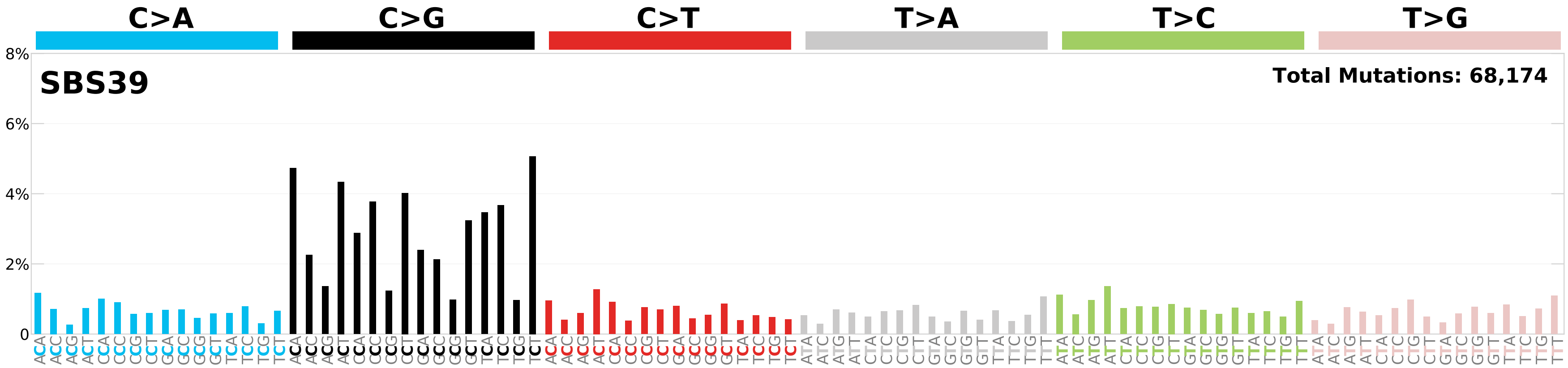

### Supplementary Fig. S9

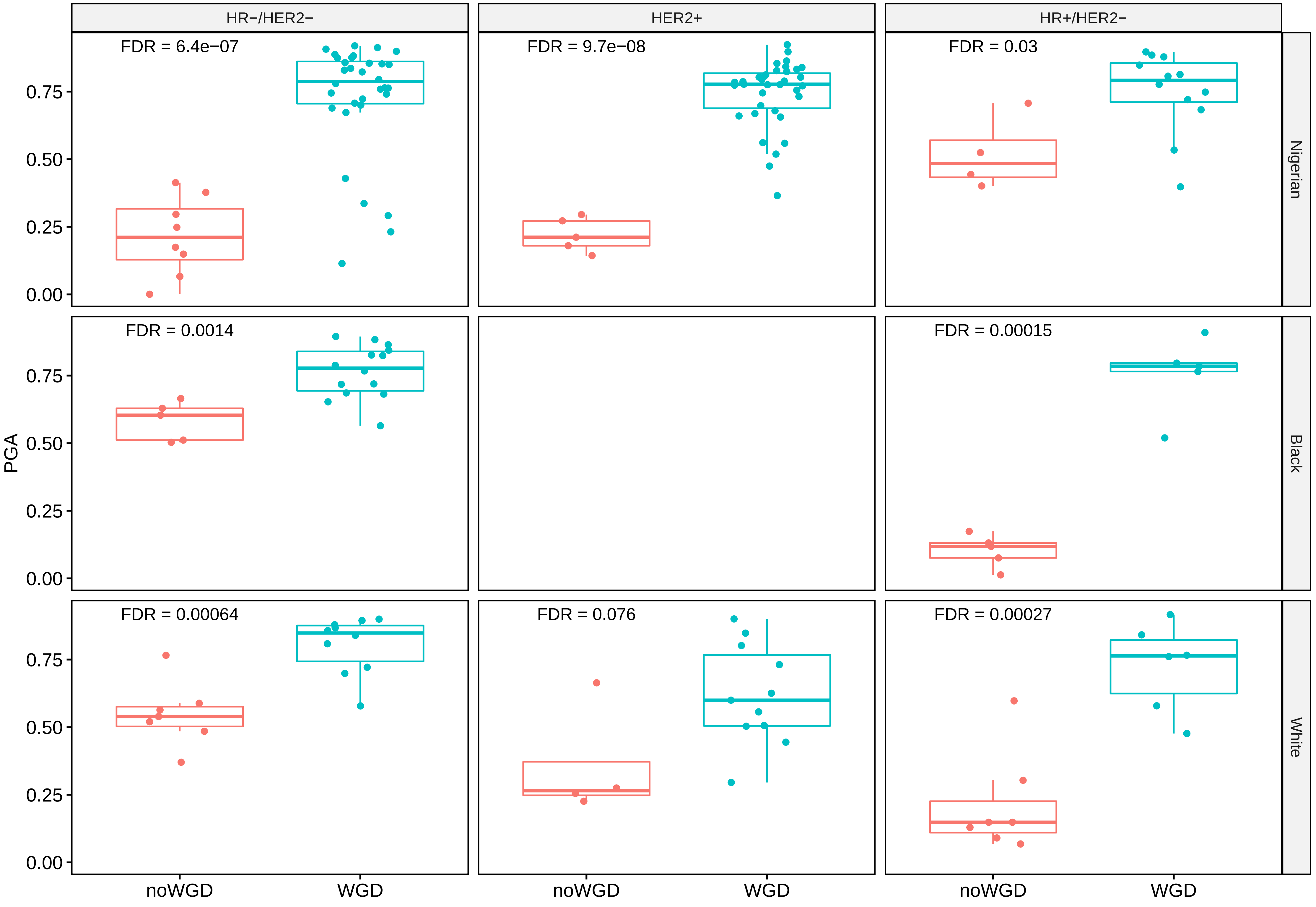

### Supplementary Fig. S10

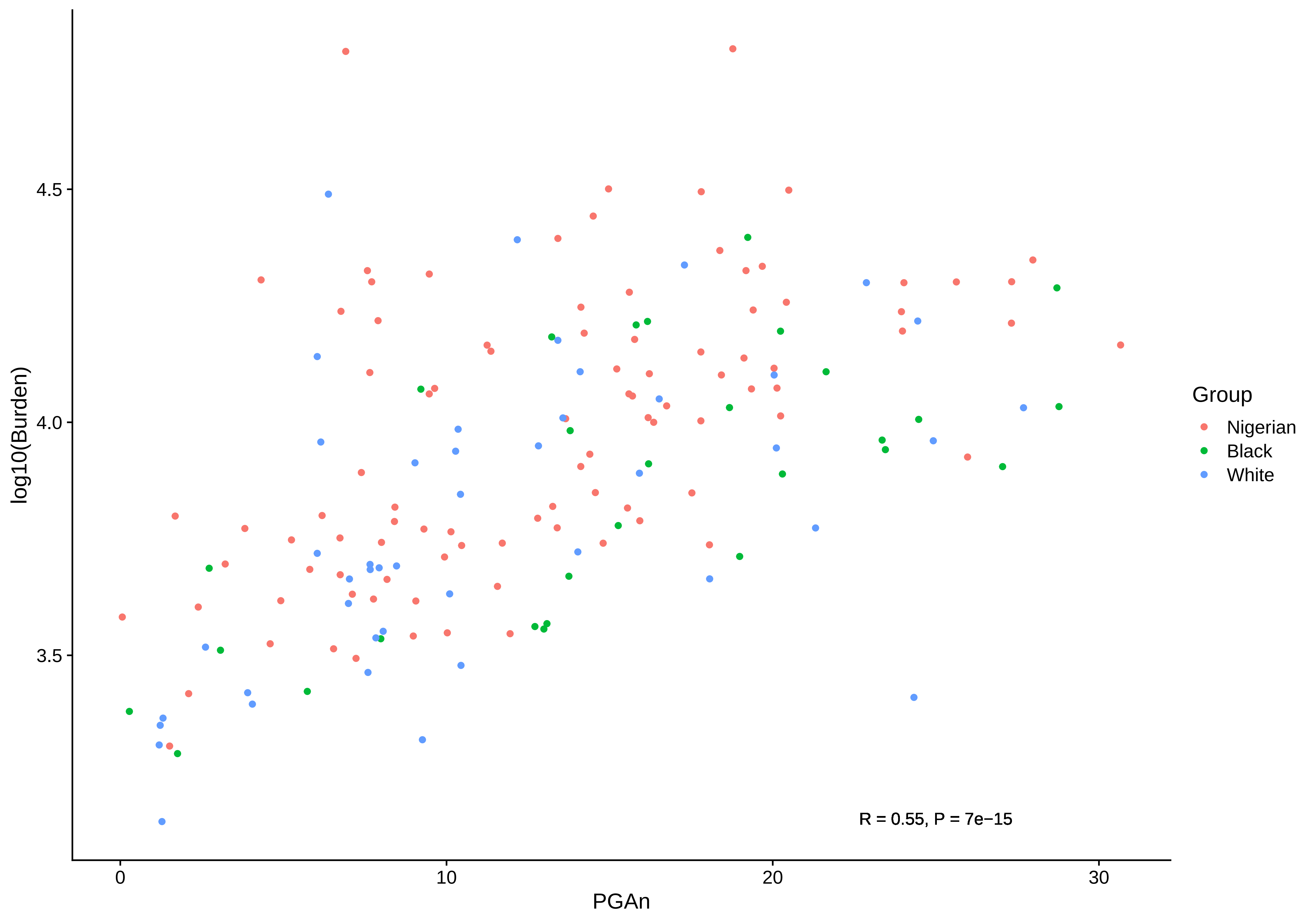

### Supplementary Fig. S11

N010842

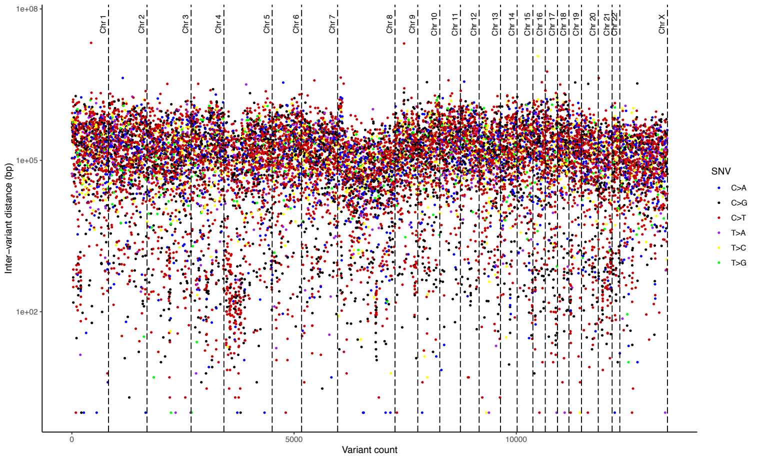


N010940

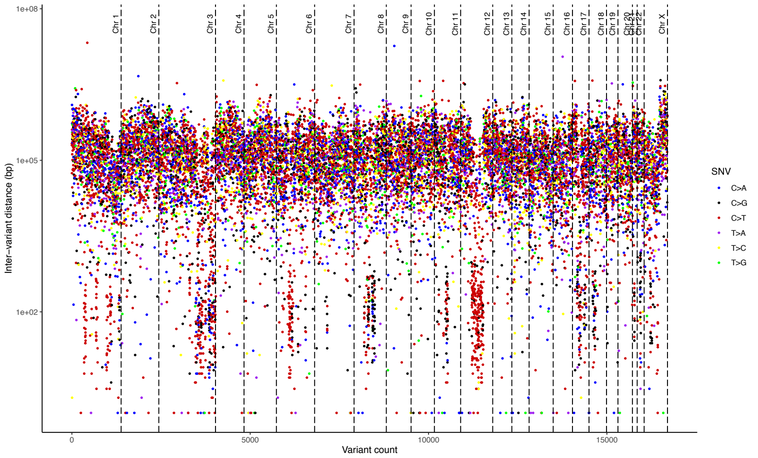


N011074

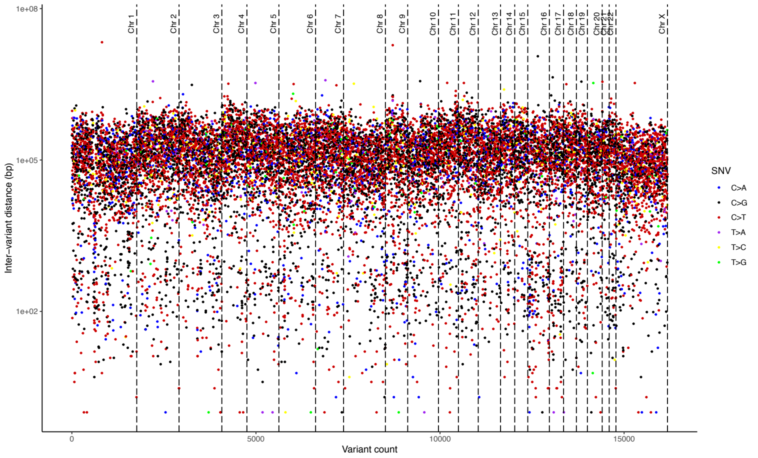


N011070

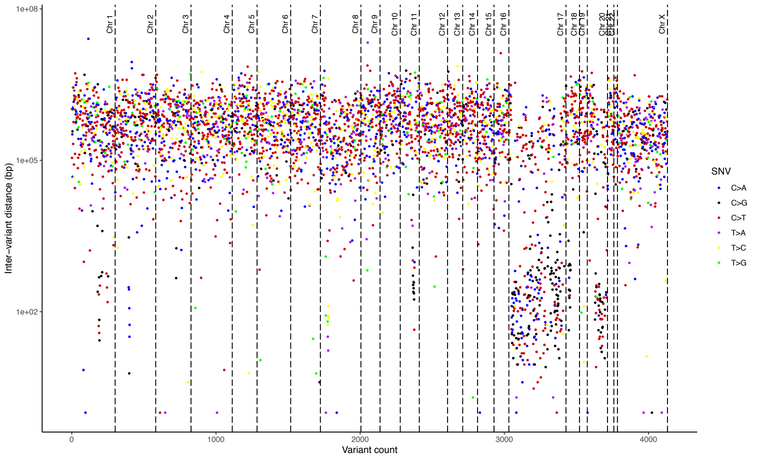

### Supplementary Fig. S12

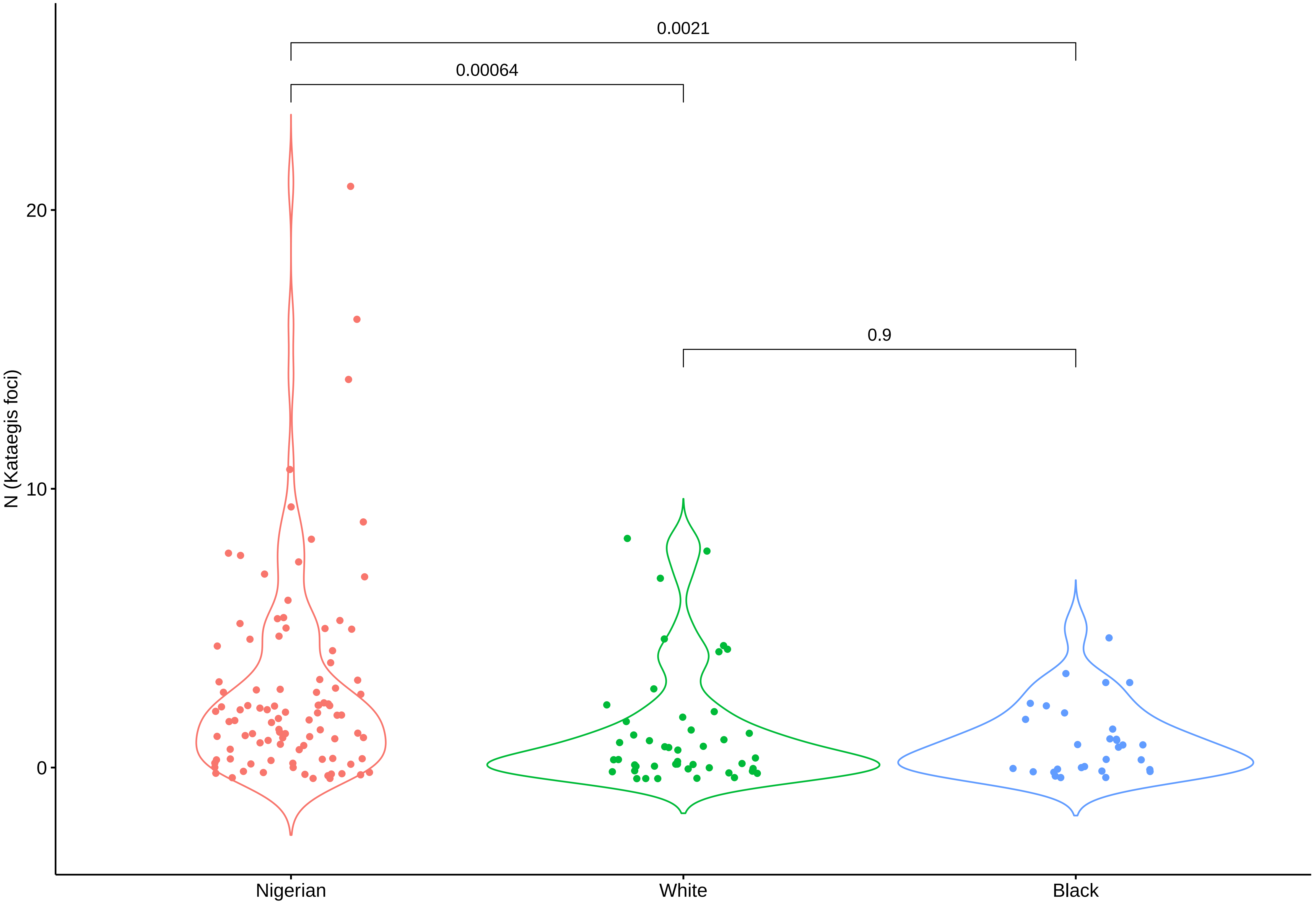

### Supplementary Fig. S13

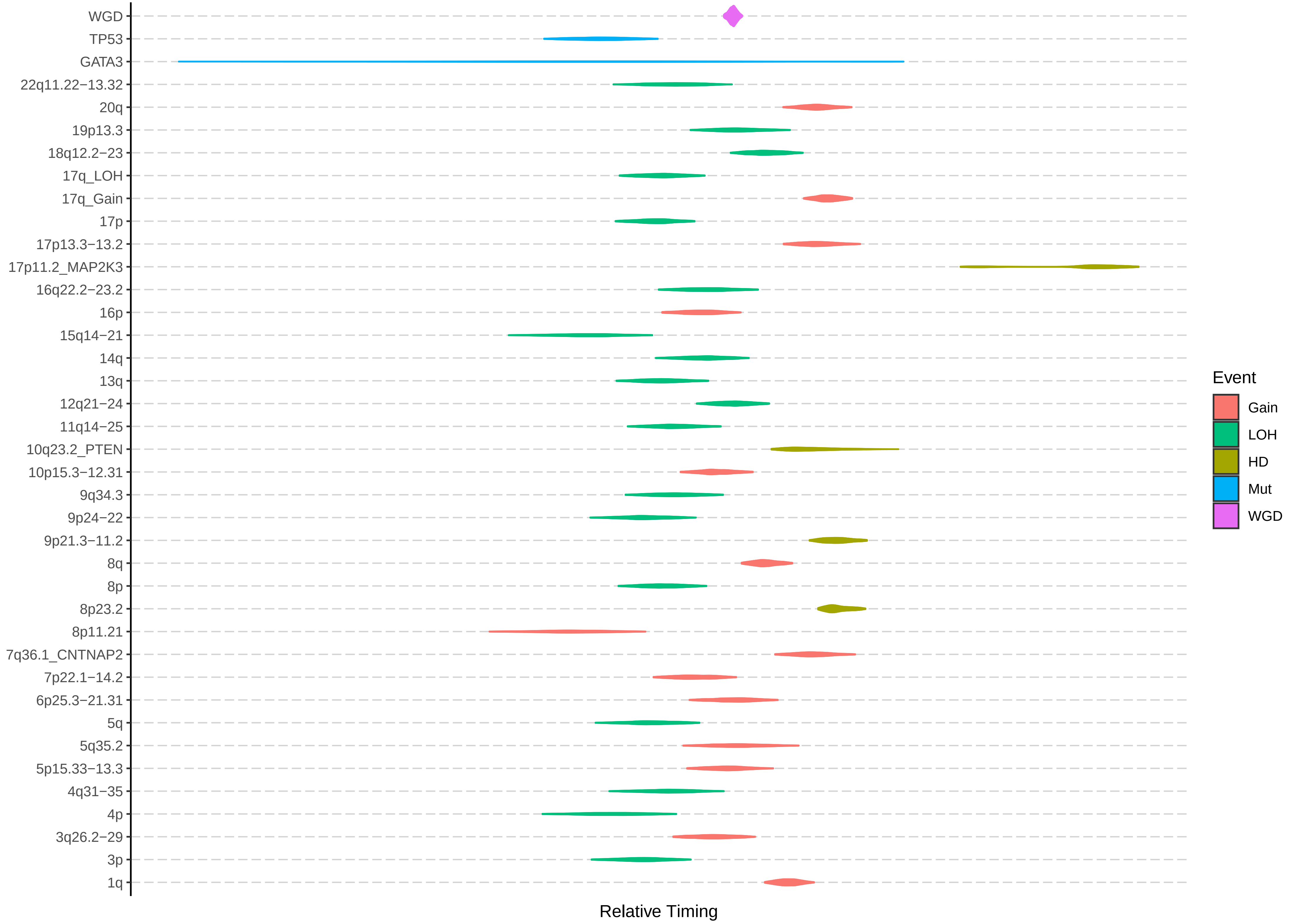

### Supplementary Fig. S14

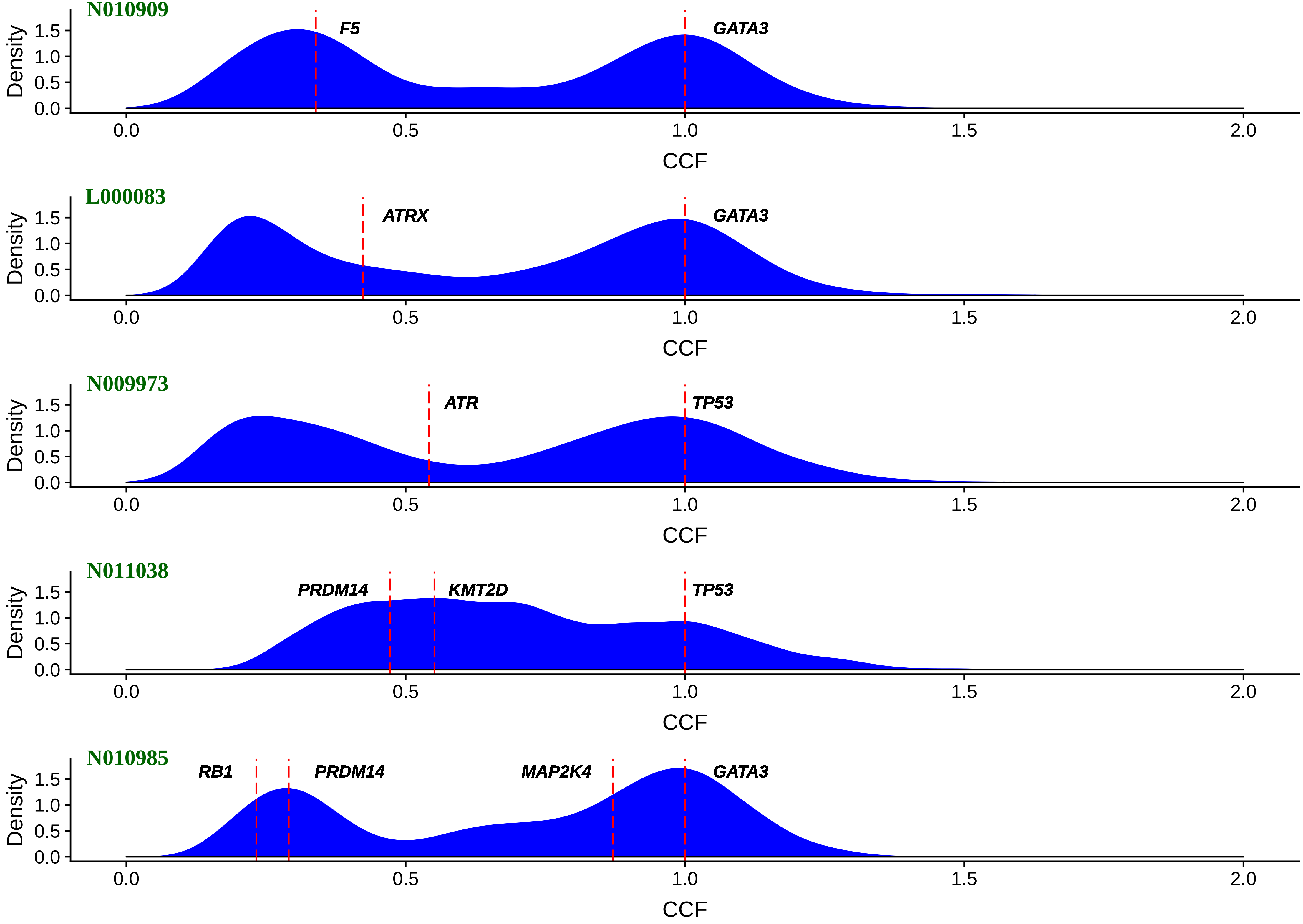

### Supplementary Fig. S15

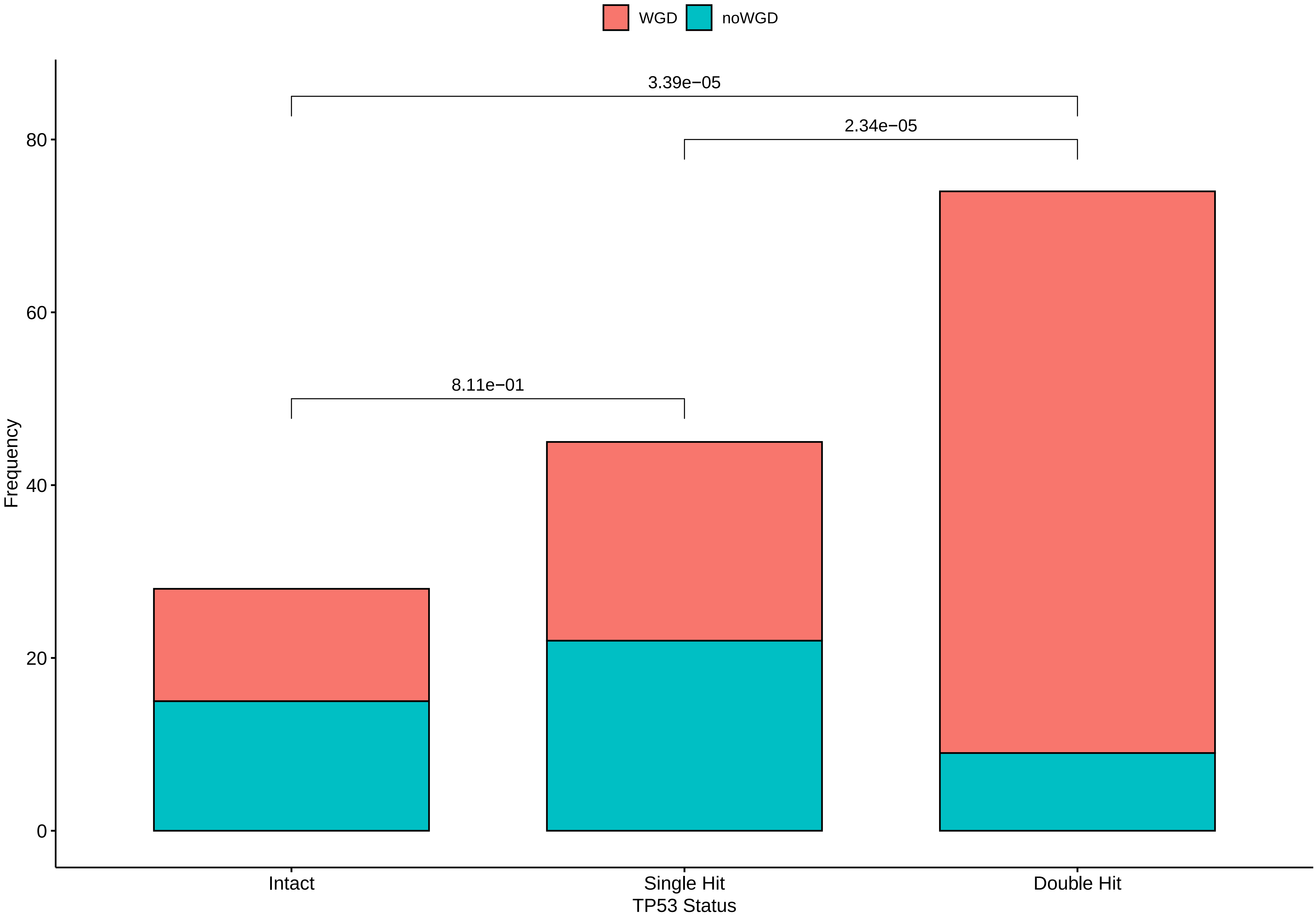

### Supplementary Fig. S16

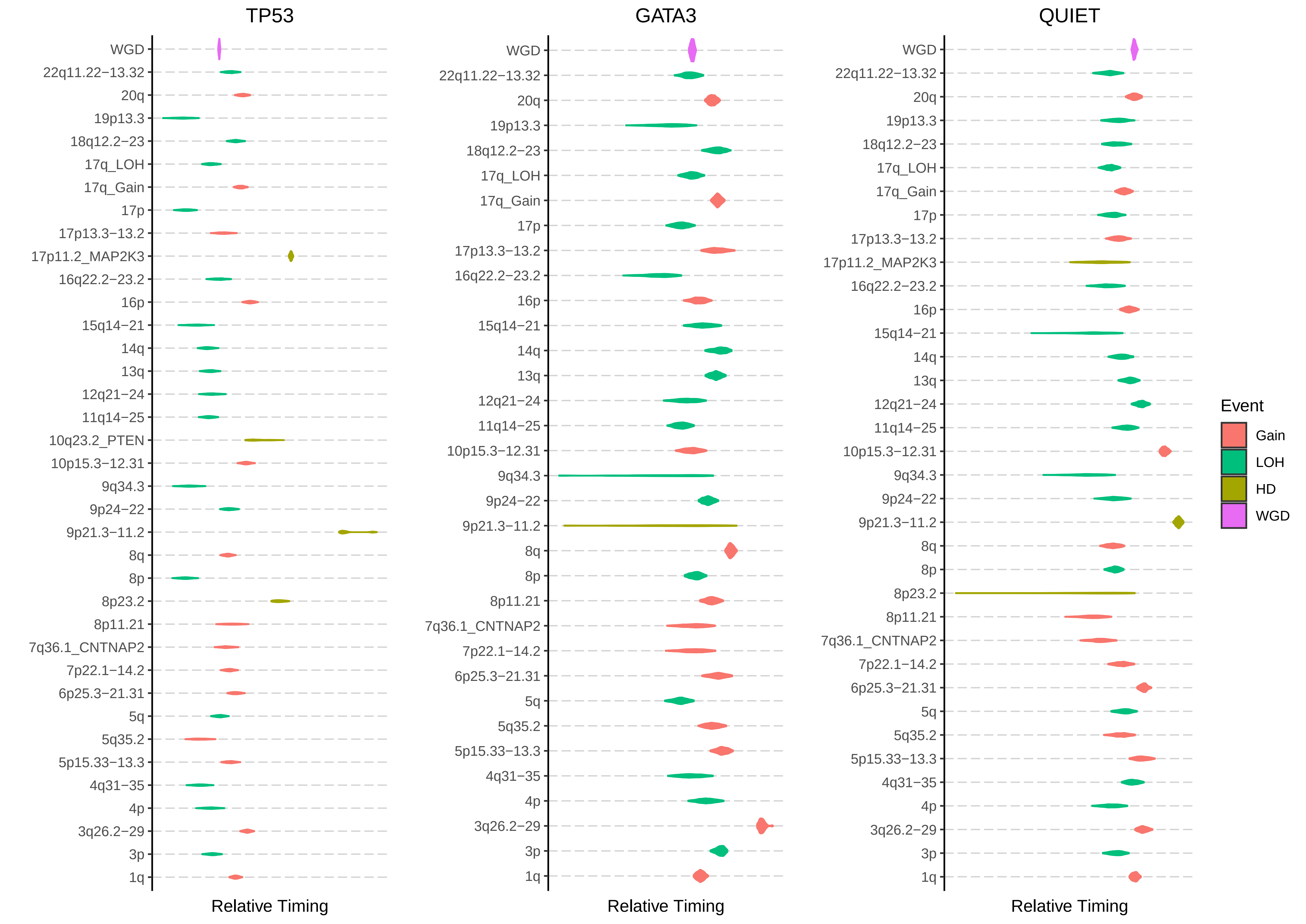

### Supplementary Fig. S17

SBS

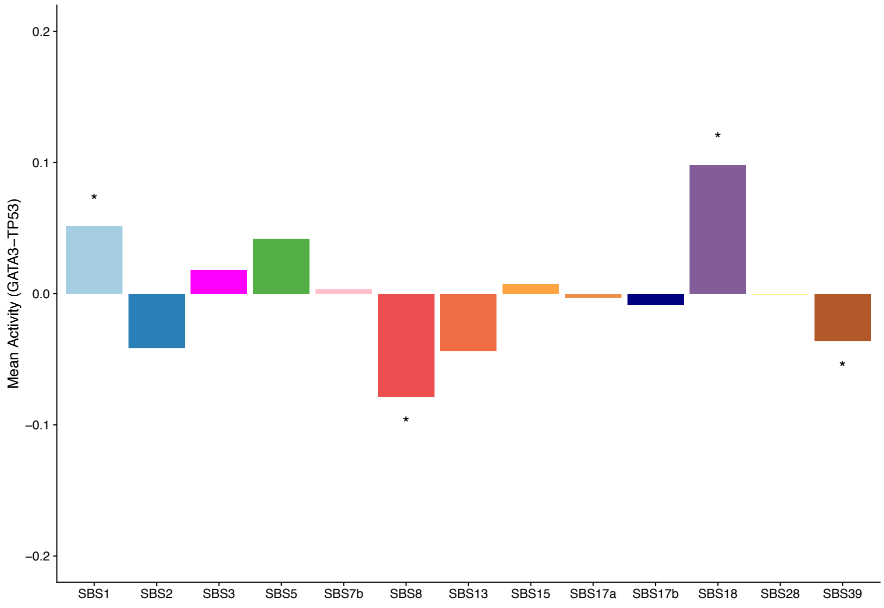


DBS

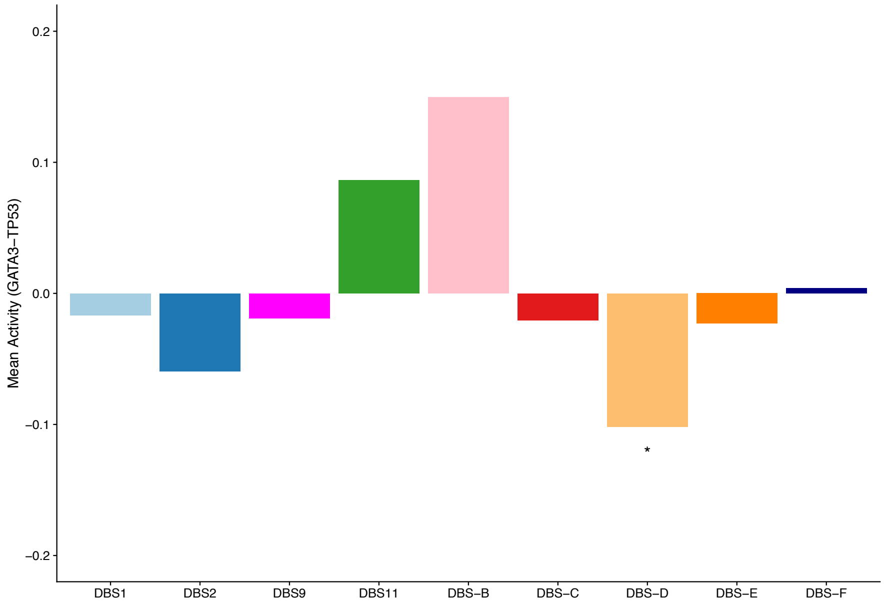


INDEL

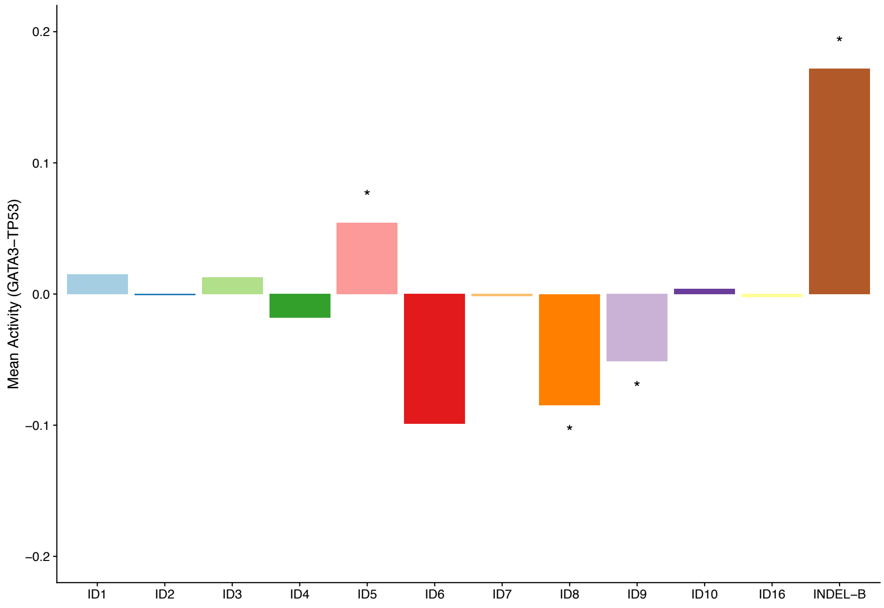

### Supplementary Fig. S18

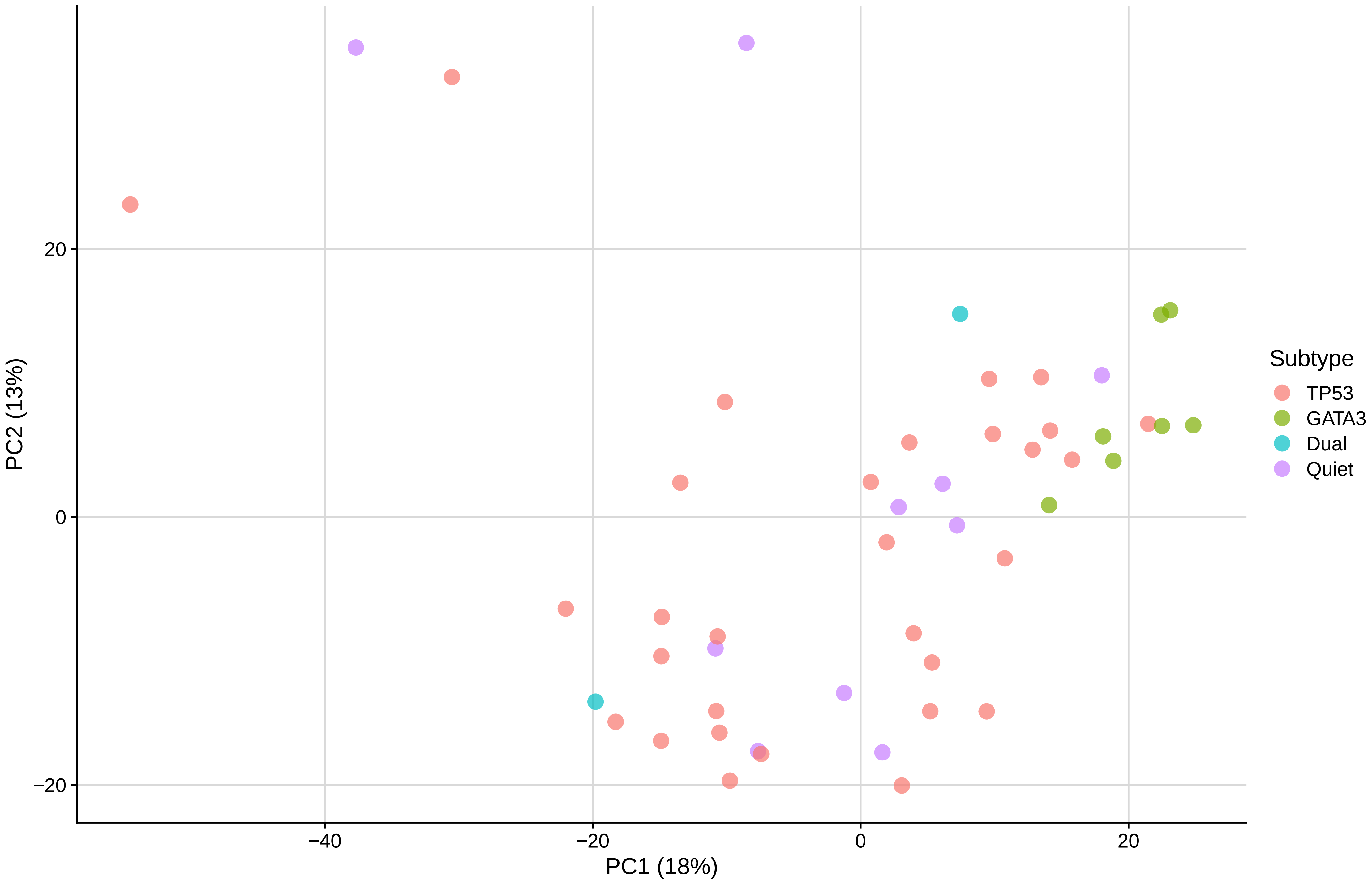

### Supplementary Fig. S19

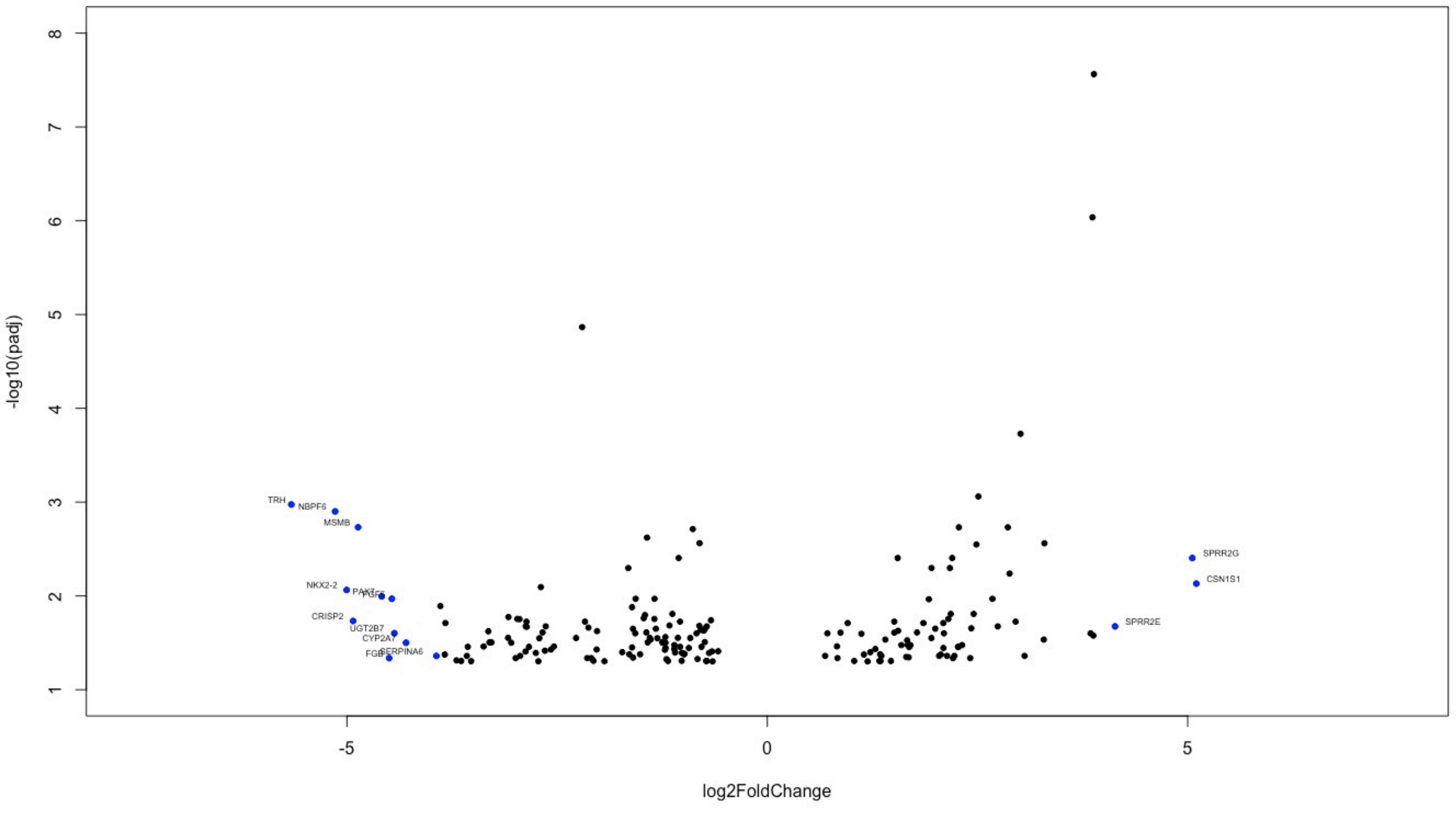
